# Structure and diversity of urinary cell-free DNA informative of host-pathogen interactions in human urinary tract infection

**DOI:** 10.1101/177402

**Authors:** Philip Burnham, Darshana Dadhania, Michael Heyang, Fanny Chen, Lars F. Westblade, Manikkam Suthanthiran, John Richard Lee, Iwijn De Vlaminck

## Abstract

Infections of the urinary tract are the most common form of infection in the human population. Here, we tested the utility of urinary cell-free DNA (cfDNA) to comprehensively monitor host and pathogen dynamics in the scope of bacterial and viral urinary tract infections. We assayed cfDNA isolated from 141 urine samples obtained from a cohort of 82 kidney transplant recipients by next-generation sequencing. We find that urinary cfDNA simultaneously informs about the composition of the bacterial and viral components of the microbiome, antimicrobial susceptibility, bacterial growth dynamics, kidney allograft injury, and the host response to infection. These different layers of information are accessible from a single assay and individually agree with corresponding clinical tests based on quantitative PCR, conventional bacterial culture, and urinalysis. In addition, cfDNA reveals the frequent occurrence of pathologies that remain undiagnosed in conventional diagnostic workups. Our work identifies urinary cfDNA as a highly versatile tool to monitor infections of the urinary tract.

## Introduction

Urinary tract infection (UTI) is one of the most common medical problems in the general population^1^. Among kidney transplant recipients, UTIs occur at an alarmingly high rate^2^. Bacterial UTI affects at least 20% of kidney transplant recipients in the first year after transplantation^3^ and at least 50% in the first three years after transplantion^4^. In addition, complications due to viral infection often occur. An estimated 5-8% of kidney transplant recipients suffer nephropathy from BK polyomavirus infection in the first three years after transplantation^5,6^. Other viruses that commonly cause complications in kidney transplantation include adenovirus, JC polyomavirus, cytomegalovirus (CMV), and parvovirus. The current gold standard for diagnosis of bacterial UTI is *in vitro* urine culture^7^. Although improved culture methods are being investigated^8,9^, bacterial culture protocols implemented in clinical practice remain limited to the detection of relatively few cultivable organisms. In addition, urinalysis is often required in conjunction with culture to make treatment decisions.

A large number of small fragments of cfDNA are present in plasma and urine^10–13^. These molecules are the debris of the genomes of dead cells from across the body and offer opportunities for precision diagnostics based on ‘*omics* principles, with applications in pregnancy, cancer and solid-organ transplantation^12,14–16^. Here, we have investigated the utility of urinary cfDNA to comprehensively monitor host and pathogen interactions that arise in the setting of viral and bacterial infections of the urinary tract. We used shotgun sequencing to assay cfDNA isolated from 141 urine samples collected from a cohort of 82 kidney transplant recipients, including patients diagnosed with bacterial UTI and BK polyomavirus nephropathy (BKVN). We implemented a single-stranded DNA (ssDNA) library preparation, optimized for the analysis of short, highly fragmented DNA^17-19^, and were able to perform sequence analyses for cfDNA isolated from relatively small volumes of urine supernatant (1 mL or less). We find that urinary cfDNA sequencing agrees in the vast majority of cases with current conventional clinical testing, while also uncovering frequent occurrence of bacteria and viruses that remain undetected in conventional diagnostic workups.

We further investigated cfDNA analysis methodologies that go beyond mere identification of microbial sequences, and that provide a deeper understanding of ongoing infections. First, we show that the pattern of cfDNA sequencing read coverage across bacterial genomes is non-uniform, with an overrepresentation of sequences at the origin of replication. A similar pattern has previously been observed in whole-genome sequencing of gut microbiota, and the disproportionate genome coverage was shown to reflect the bacterial growth rate, where an overrepresentation of genomic coverage at the replication origin signals faster population growth^20^. We show that measuring the bacterial population growth rate from urinary cfDNA can be used to inform diagnosis of UTI. Second, we mined cfDNA for antimicrobial resistance (AR) genes and show that AR gene profiling can be used to evaluate antimicrobial resistance. Furthermore, we demonstrate that cfDNA informs about the host response to infection on both a cellular and tissue level. Recent reports have demonstrated that cfDNA in plasma comprises the footprints of DNA-binding proteins and nucleosomes^21,22^. We find that nucleosome structures within transcription regulatory elements are preserved in urinary cfDNA, as was previously described for plasma cfDNA^22^. The occupancy of nucleosomes in gene regions flanking the transcription start site is notably reduced for transcribed genes, consistent with known chromatin alterations in gene promoters during transcription, thus providing a measure of gene expression. Furthermore, we observe that the relative proportion of kidney donor specific cfDNA signals graft tissue injury in the setting of viral infection and host immune cell activation in the scope of bacterial infection. Finally, we report that the graft is the predominant source of mitochondrial cfDNA in the urine of kidney transplant recipients with BKVN.

Collectively, this study supports the utility of shotgun sequencing of urinary cfDNA as a comprehensive tool for monitoring patient health and studying host-pathogen interactions.

## Results

### Biophysical properties of urinary cfDNA

Urinary cfDNA is comprised of chromosomal, mitochondrial, and microbial cfDNA released from host cells and microbes in the urinary tract, and of plasma-derived cfDNA that passes from blood into urine^23^. Urine can be collected non-invasively in large volumes, and therefore represents an attractive target for diagnostic assays. Compared to plasma DNA, relatively few studies have examined the properties and diagnostic potential of urinary cfDNA. The urinary environment degrades nucleic acids more rapidly than plasma resulting in fewer DNA fragments that are shorter^24^. Consequently, sequence analyses of urinary cfDNA have to date required relatively large (> 10 mL) volumes of urine^13,25^. Here, we applied a single-stranded library preparation technique that employs ssDNA adapters and bead ligation to create diverse sequencing libraries that capture short, highly degraded cfDNA^18,19^ (Fig. 1A). We find that single-stranded library preparation enables sequence analyses of urinary cfDNA from just one milliliter of urine supernatant. We assayed 141 urine samples collected from kidney transplant recipients, including subjects diagnosed with bacterial UTI and polyomavirus nephropathy (overview of post-transplant dates and categories depicted in Fig. 1B, see Methods for details). We obtained 43.5 +/- 17.3 million paired-end reads per sample, yielding a per-base human genome coverage of 0.49x +/- 0.24x. Many fragments derived from microbiota; for example, for patients diagnosed with bacterial UTI, bacterial cfDNA accounted for up to 34.65% of the raw sequencing reads and in cases of BKVN, BK polyomavirus cfDNA accounted for up to 10.27% of raw sequencing reads. To account for technical variability and sources of environmental contamination during extraction and library preparation, a known-template control sample was included in every sample batch and sequenced (see Methods).

**Figure 1.**
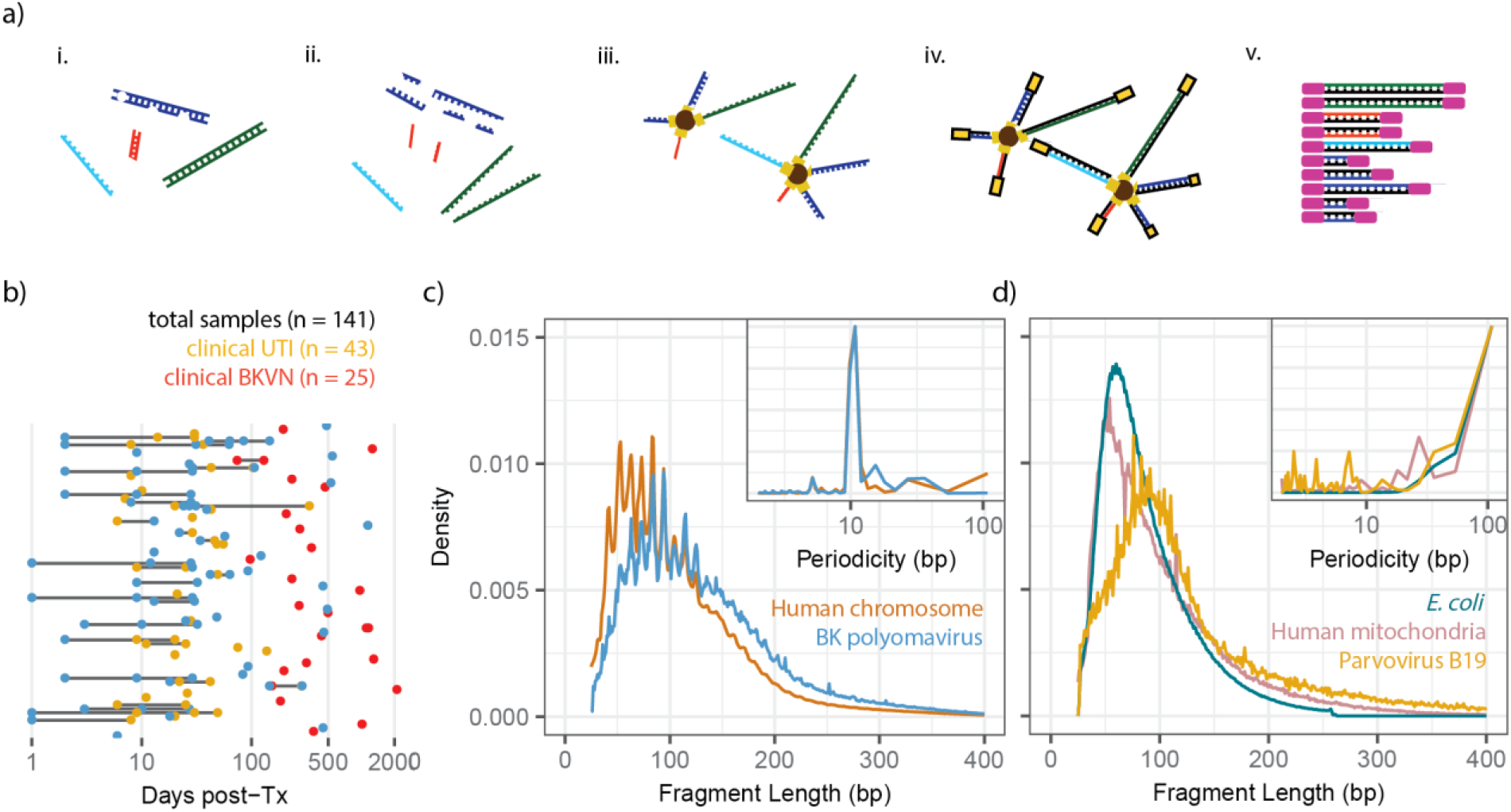
Shotgun sequencing assay and biophysical properties of urinary cfDNA. **(a)** Schematic representation of the ssDNA library preparation protocol used for shotgun sequencing of urinary cfDNA^19^. Key steps include: *i*) cfDNA isolation, *ii*) DNA denaturation, *iii*) ssDNA adapter ligation, *iv*) extension and double-stranded DNA adapter ligation, *v*) and PCR. **(b)** Overview of post-transplant sample collection dates (color indicates pathology, bars connect samples from same patients). **(c-d)** Fragment length density plot measured by paired-end sequencing for different cfDNA types: (**c**) chromosomal and polyomavirus cfDNA from representative samples, and (**d**) *E. coli*, parvovirus, and mitochondrial cfDNA from representative samples. Fourier analysis reveals a 10.4 bp periodicity in the fragment length profiles of chromosomal and BK polyomavirus cfDNA but not in *E. coli*, parvovirus B19, and mitochondrial cfDNA (insets). See supplemental table “Clinical Data”.

We analyzed the fragment length profiles of urinary cfDNA at single nucleotide resolution using paired-end read mapping^10^. This analysis confirmed previous observations of the highly fragmented nature of urinary cfDNA compared to plasma cfDNA^25^ (Fig. 1C). We observed a 10.4 bp periodicity in the fragment length profile of chromosomal cfDNA (Fourier analysis, Fig. 1C, inset), consistent with the periodicity of DNA-histone contacts in nucleosomes^26^. Polyomavirus is known to hijack histones of infected host cells, and to form mini chromosomes after infection^27^. The periodicity in the fragment length profiles of BK polyomavirus cfDNA in urine reflect this biology (Fig. 1C). We did not observe a similar nucleosomal footprint for bacterial and mitochondrial cfDNA, and cfDNA arising from parvovirus B19, which is expected given the non-nucleosomal compaction of the genomes that contribute these cfDNA types (Fig. 1D).

### Infectome screening

We assessed the presence of cfDNA from bacterial and viral pathogens reported by conventional diagnostic assays. We used previously described bioinformatics approaches to quantify non-human cfDNA^28^. Briefly, human sequences were identified by alignment to a human reference genome and removed. Remaining sequences were BLASTed against a database of microbial reference genomes. We estimated the relative genomic representation of different species using GRAMMy^29^. To directly compare the measured microbial abundance across samples and species, we computed the representation of microbial genome copies relative to the representation of the human genome, and expressed this quantity as relative genome equivalents (RGE).

We detected a very high load of BK polyomavirus cfDNA in all 25 samples collected from 23 patients diagnosed with BKVN by needle biopsy (mean 1.49 +/- 1.08 x10^5^ RGE, Fig. 2A), but not in samples from 11 patients that were BKVN negative per biopsy (all below detection limit). The BK polyomavirus cfDNA abundance (RGE) correlated with a matched urine cell pellet BKV VP1 mRNA copy measurement that we previously validated as a noninvasive marker for BKVN^30,31^ (Spearman: *ρ* = 0.73, *ρ* = 7.1 x10^−7^).

**Figure 2.**
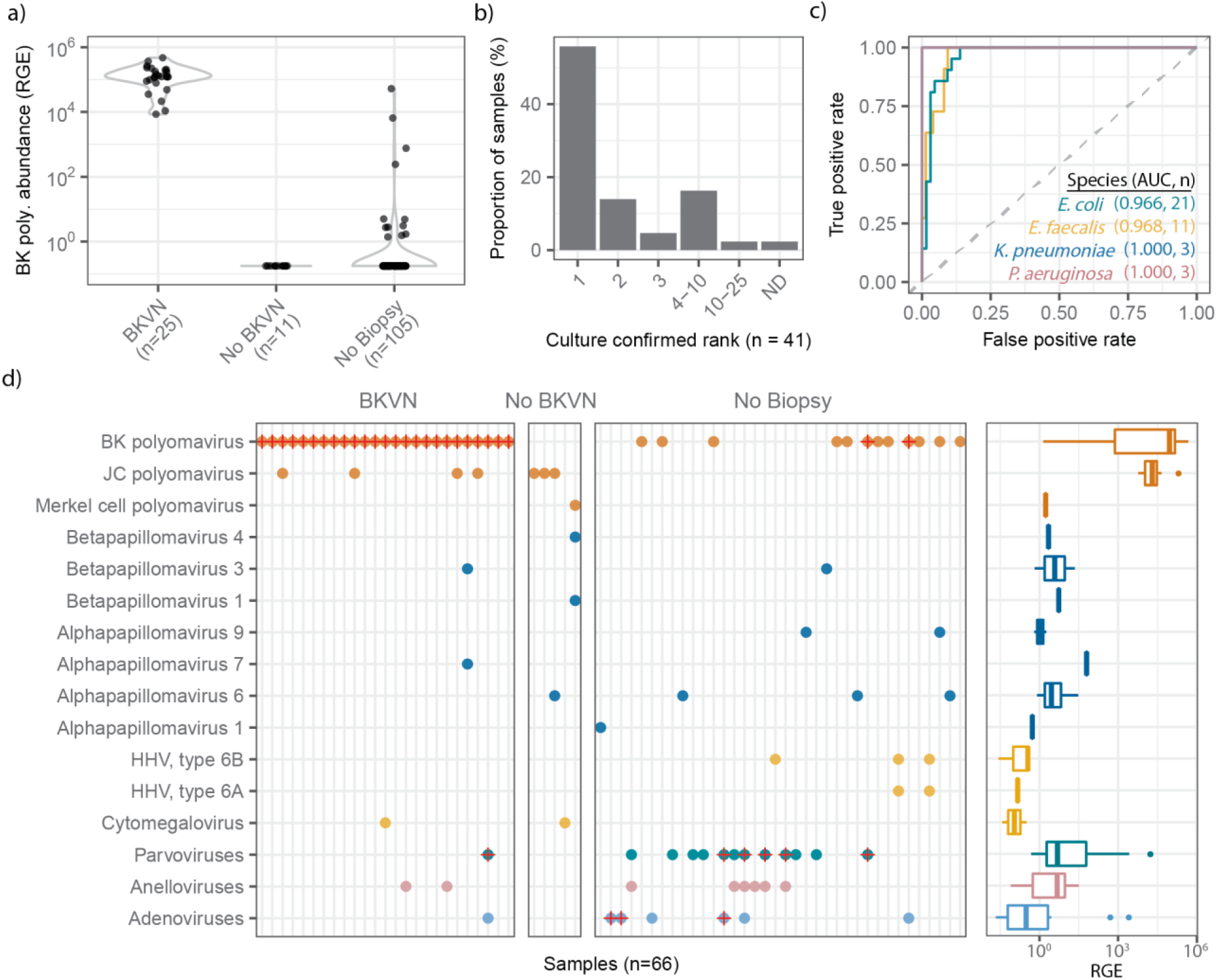
Urinary cfDNA infectome screening. **(a)** Violin plots of BK polyomavirus cfDNA sequence abundance (RGE) for patients with and without BKVN and untested patients. **(b)** cfDNA rank order abundance for clinically reported uropathogens. In 60% of samples the bacterial organism detected in culture was the most abundant component of the cfDNA urinary microbiome. In one sample, the clinically reported agent was not detected (ND). **(c)** Receiver operating characteristic analysis of the performance of urinary cfDNA in identifying UTIs due to common bacterial uropathogens (86 urine samples, AUC = area under the curve, n = number of positive cultures, see Fig. S1 for individual Receiver Operating Characteristic curves for these and four additional species) **(d)** cfDNA reveals frequent occurrence of viruses that are potentially clinically relevant (left panel); red crosses identify samples belonging to patients who developed an infection of the corresponding viral group. Right panel shows boxplots of the viral cfDNA abundance across all samples (right panel). Coloring of points and boxplots by viral taxonomic group; see supplemental table “Fig2A-D”.

We quantified bacterial urinary cfDNA in 43 urine samples from 31 patients who had a corresponding same day positive culture. For 41 of the 43 clinically positive urine specimens, a particular bacterial species was reported by conventional culture. In 40 of these 41 samples, sequencing of urinary cfDNA detected the clinically reported suspected uropathogen to the species level (Fig. 2B). For a single sample, urinary cfDNA did not correlate with the bacterial culture. *Raoultella ornithinolytica* was isolated in culture, but not detected in cfDNA (see Methods for a detailed discussion of this discordant readout). For two clinically positive samples, the primary suspected causative agent was identified to the genus level (*Staphylococcus [reported as coagulase-negative Staphylococcus species]* and *Streptococcus*) by conventional culture. In both these cases the primary agent was detected as the most prevalent within the sample. We examined five samples from patients with polymicrobial infection (defined as at least two bacterial species detected at > 10,000 CFU/mL in culture). For four out of five of these cases, we observed both species among the ten most abundant species (see Supplementary data table for these cases). In one sample, the secondary bacterial agent, coagulase-negative *Staphylococcus* species (CoNS), was not detected.

To further test the performance of urinary cfDNA to identify specific bacteria, we compared the relative abundance of bacterial cfDNA for patients diagnosed with bacterial infection (48 bacterial isolates identified from 43 conventional cultures), to the relative genomic abundance measured for 43 negative urine cultures (defined as < 10,000 CFU/mL), (Fig. 2C and Fig. S1). We find very good agreement between urinary cfDNA and culture based isolation of *Enterococcus faecalis* (number of matched positive cultures, n = 11, Area Under the Curve, AUC = 0.97), *Enterococcus faecium* (n = 2, AUC = 0.98), *Escherichia coli* (n = 21, AUC = 0.97), *Klebsiella pneumonia* (n = 3, AUC = 1.00), *Pseudomonas aeruginosa* (n = 3, AUC = 1.00), *Klebsiella oxytoca* (n = 1, AUC = 1), CoNS (n = 4, AUC = 0.78), and viridans group streptococci (n = 1, AUC = 0.98).

In only 60% of examined samples (26/43 UTI cases), we found that the uropathogen identified by culture was the most prevalent pathogen in the sample (Fig. 2B). Whereas bacterial culture is skewed towards species that are readily isolated on routine bacteriological media employed for urine culture, cfDNA sequence analyses potentially permit the identification of a broader spectrum of bacterial species. To evaluate this concept further, we assayed two samples collected from a patient diagnosed with *Haemophilus influenzae* bacteruria. *H. influenzae* is a very uncommon uropathogen that does not routinely grown on media employed for conventional urine culture (tryptic soy agar with sheep blood and MacConkey agar)^32^. Repeated cultures for this patient were negative, but given a urinalysis suggestive of a UTI and given that the patient developed *H. influenzae* bacteremia, the original urine specimen collected at presentation was replated on chocolate agar, upon which *H. influenzae* was isolated. In the sample taken at the time of presentation, which was cultured, and also a sample taken four days after presentation, we observed a high abundance of *H. influenzae* cfDNA (0.037 RGE and 0.41 RGE respectively). This case supports the utility of urinary cfDNA to identify infections where conventional culture fails.

### Profiling the Urinary Microbiome

The urinary tract was long regarded as sterile but recent studies have revealed that the urinary tract often harbors unique microbiota^9,33,34^. We have examined the composition of the urinary microbiome by urinary cfDNA profiling in the absence of bacterial UTI (Fig. S2). We find that the species level abundance and the species level diversity of the bacteriome are a function of the transplant recipient gender but not the donor gender. On average, we observed two to three orders of magnitude more cfDNA from *Gardnerella* (6125x), *Ureaplasma* (1686x), and *Lactobacillus* (321x) species across female transplant recipients who did not have a UTI at time of sampling compared to male recipients who did not have UTI; these bacterial genera are well characterized as members of the healthy female vaginal microbiome^35^. We examined the relationship between urine collection methods and the abundance and diversity of the bacteriome and find a notably reduced bacterial load for samples collected by Foley catheter (samples collected within four days after transplant) versus clean catch urine samples. cfDNA may be an ideal tool to study the urinary microbiome, but such future studies need to account for effects of gender and sample collection approaches.

### Broad screening for viruses via cfDNA

We next screened for the occurrence of cfDNA derived from viruses. Nearly half of the samples (66/141) had detectable levels of cfDNA derived from eukaryotic viruses that are potentially clinically relevant. Figure 2D highlights the frequent occurrence of JC polyomavirus, parvovirus B19, Merkel cell polyomavirus, cytomegalovirus (CMV), human herpesvirus 6A, human herpesvirus 6B, and various known oncoviruses across different patient groups. In several samples, we detected cfDNA from multiple polyomavirus species concurrently (JC polyomavirus or BK polyomavirus). To shed light on the potential clinical utility of broad screening for viruses via cfDNA, we assayed serial urine from three patients diagnosed with viral infections that are relatively uncommon in kidney transplant recipients and consequently not routinely screened for in our patient cohorts. In samples from two patients with clinically diagnosed parvovirus B19 infection, we detected urinary cfDNA from parvovirus B19 up to 8 days prior to the clinical diagnosis in one subject and urinary cfDNA from parvovirus B19 up to 80 days before diagnosis and up to 25 days after diagnosis in another subject. In the former subject, we observed a high abundance of both BK polyomavirus (3.54 x10^4^ RGE) and parvovirus B19 (2.48 x 10^4^ RGE), which correlated with positive results of individual viral-specific PCR tests for BK polyomavirus and parvovirus B19 performed in the clinic. For a third patient, we observed a high abundance of human adenovirus B DNA, in samples obtained up to 15 days before (2.52 x 10^3^ RGE), and 9 days after (5.08 x10^2^ RGE) adenovirus infection diagnosis. These data support the utility of urinary cfDNA sequencing for the detection of both common and uncommon viral agents.

### Quantifying bacterial growth rates

Conventional metagenomic sequencing can provide a snapshot of the microbiome, yet does not inform about microbial life cycles or growth dynamics. In a recent study, Korem *et al.* reported that the pattern of metagenomic sequencing read coverage across a microbial genome can be used to quantify microbial genome replication rates for microbes in complex communities^20^. Here, we tested whether this concept can be used to estimate bacterial population growth from measurements of cfDNA. Figure 3A shows the urinary cfDNA sequence coverage for four bacterial species, *E. coli*, *K. pneumoniae*, *Gardnerellla vaginalis* and *Cutibacterium acnes*. For two patients diagnosed with *E. coli* and *K. pneumoniae* UTI (Fig. 3A), the *E. coli* and *K. pneumoniae* genome coverage was highly non-uniform, with an overrepresentation of sequences at the origin of replication and an underrepresentation of sequences at the replication terminus. The shape of the *E. coli* and *K. pneumoniae* genome coverage is a result of bi-directional replication from a single origin of replication. The skew in genome coverage reflects the bacterial population growth rate, where a stronger skew signals faster population growth^36^. The genome coverage of a typically commensal bacterial species, *G. vaginalis*, exhibited non-uniform genome coverage (Fig. 3A), similar to the above uropathogens but less pronounced. *C. acnes* has been recognized as a common skin and lab contaminant^37^. The genome coverage for *C. acnes*, was highly uniform, indicative of slow or no growth (aggregate across 99 samples, Fig. 3A).

**Figure 3.**
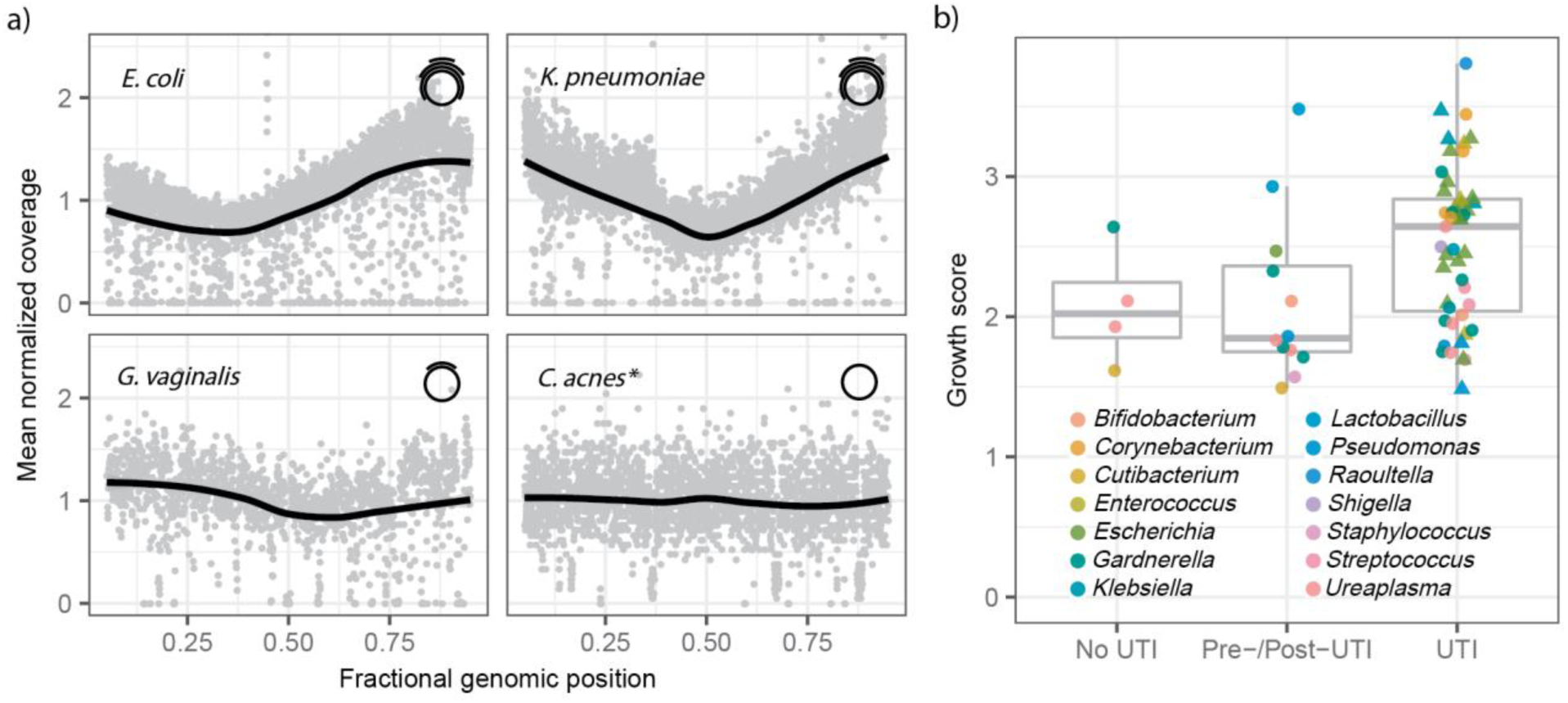
Estimating bacterial population growth rates from urinary cfDNA. **(a)** Normalized bacterial genome coverage for four representative bacterial species. The coverage was binned in 1 kbp tiles and normalized. Each panel represents a single sample (see Supplementary data table), with the exception of *C. acnes* (*) for which the coverage was aggregated across 99 samples (solid line is a LOESS filter smoothing curve, span = 0.70). The non-uniform genome coverage for *E. coli* and *K. pneumoniae*, with an overrepresentation of sequences at the origin of replication, is a result of bidirectional replication from a single origin of replication. The initial and final 5% of the genome is removed for display. **(b)** The skew in genome coverage reflects the bacterial growth rate, where a stronger skew signals faster growth^36^. Box plots of growth rates for species in twelve genera grouped by patient groups (at least 2500 alignments, 41 samples, see methods for definition of Pre/Post-UTI). Each point indicates a bacterial species in a sample. Triangles indicate culture-confirmed bacteria by genus. Boxplot features describes in Methods. See supplemental table “Fig3A-B”.

We asked whether this measure of bacterial growth can be used to inform UTI diagnosis. We calculated an index of replication based on the shape of the sequencing coverage using methods described previously^36^. We used BLAST to identify abundant bacterial strains and then re-aligned all sequences with BWA to a curated list of bacterial species. Samples for which the genome coverage was too sparse were excluded from this analysis (see Methods). Figure 3B compares the index of replication for bacteria in samples from patients diagnosed with UTI, to bacteria in samples from patients with negative cultures and samples collected from patients before and after UTI diagnosis. Species categorized in the UTI group had markedly greater growth rates, than those in the no UTI and pre-/post-UTI groups (two-tailed Wilcox rank sum test, *p* = 9.0 x 10^−3^).

### Antimicrobial resistome profiling

For 42 of 43 samples collected from patients with clinically confirmed UTIs, we determined the relative abundance of genes conferring resistance to several classes of antimicrobials (a single sample, for which no AR gene fragments were observed, was excluded from this analysis). We used blastp to align non-human sequences against known AR genes and mutations^38^. AR gene sequences were aggregated and called against the non-redundant Comprehensive Antibiotic Resistance Database that indicates the drug resistance conferred by the given gene.

We compared the results of phenotypic antimicrobial susceptibility testing (see Methods) to the resistance profiles determined by sequencing. For most samples, there was a high diversity in alignments with highly abundant resistance classes including resistance to macrolides, aminoglycosides, and beta-lactams (Fig. 4). We studied vancomycin-resistant *Enterococcus* (VRE) infections, which often lead to complications after transplantation, in depth. Resistance to vancomycin was clinically assessed via measurement of the minimum inhibitory concentration value using broth microdilution (MicroScan, Beckman Coulter, Inc) or gradient diffusion (Etest®, bioMérieux, Inc). We detected fragments of genes conferring resistance to the glycopeptide antibiotic class, of which vancomycin is a member, for all VRE positive samples (n = 4). Moreover, for samples with *Enterococcus* that tested as vancomycin susceptible (n = 7), we did not detect fragments of glycopeptide class resistance genes (Fig. 4). These data indicate significant potential to predict antimicrobial susceptibility from measurements of urinary cfDNA.

**Figure 4.**
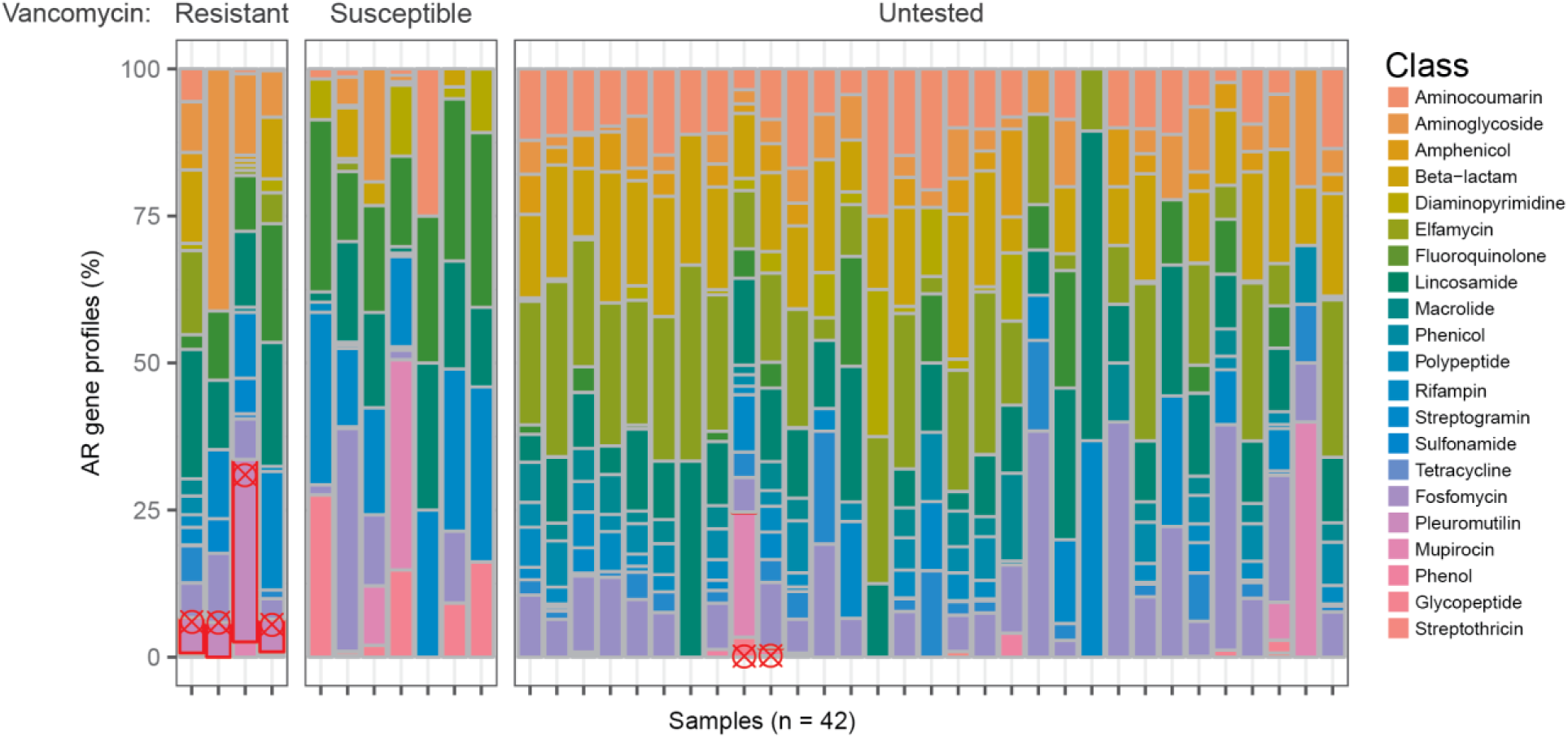
cfDNA based antimicrobial resistome profiling. For 42 samples from patients with clinically confirmed UTI, AR gene profiling reveals the presence of genes conferring resistance to various drug classes. The data is organized in three sample groups: samples from patients with vancomycin resistant *Enterococcus* (Resistant), samples from patients with vancomycin susceptible *Enterococcus* (Susceptible), and samples from patients for which vancomycin resistance testing was not performed (Untested). Samples in which fragments of genes that confer resistance to glycopeptide class antibiotics (including vancomycin, red outlines) were detected, are marked by red crosshairs. See supplemental table “Fig4”.

### Host response to infection

We next examined the host response to viral and bacterial infections. Recent work has identified transplant donor specific cfDNA in plasma as a marker of graft injury in heart, lung, liver and kidney transplantation^15,28,39,40^. Here, we quantified donor specific cfDNA in urine for sex-mismatched donor recipient pairs by counting Y chromosome derived cfDNA (Fig. 5A, Methods). We observed elevated levels of donor cfDNA in the urine of patients diagnosed with BKVN (mean proportion of donor DNA 65.1%, n=12) compared to the urine of patients who had normal biopsies (no BKVN, mean 42.2%, n=5) and samples from patients who did not develop a clinical UTI in the first three months of transplantation (mean 25.5%, n=11, samples collected within five days after transplant excluded). The release of donor DNA reflects severe cellular and tissue injury in the graft, a hallmark of BKVN. In contrast to patients with BKVN, patients diagnosed with bacterial UTI had lower proportions of donor DNA as compared to stable individuals. This is likely explained by an elevated number of recipient immune cells in the urinary tract following immune activation. Indeed, comparison to clinical urinalysis indicates that the donor fraction decreases with increasing white blood cell count (WBC, per high power field, HPF, 400 x microscope magnification, inset Fig. 5A, Spearman: *ρ* = −0.57, *p* = 1.3 x 10^−4^). Furthermore, clinical cases of pyuria – defined as greater than ten WBC per HPF^41^, had a lower donor fraction than those without (two-tailed Wilcox test, *p* = 8.0 x 10^−4^). In addition, we found that the level of donor DNA in the first few days after transplant was elevated, consistent with early graft injury. We tracked the relative and absolute abundance of donor specific urinary cfDNA in the first few days after transplantation for a small subset of subjects (n=5). The initial elevated level of donor DNA quickly decayed to a lower baseline level (Fig. 5B), in line with previous observations in heart and lung transplantation^15,42^.

**Figure 5.**
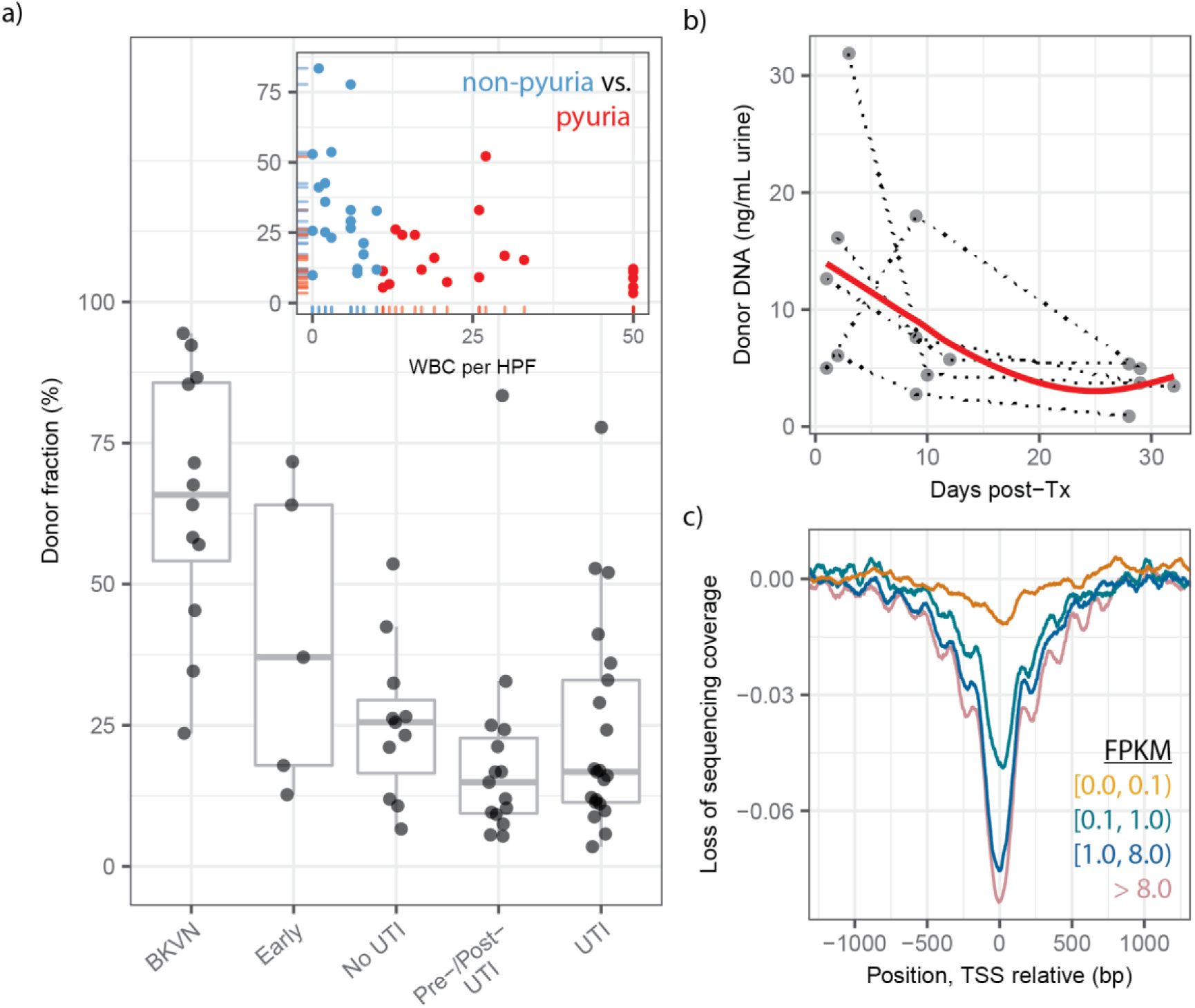
Quantifying the host response to infection from urinary cfDNA. **(a)** Proportion of donor-specific cfDNA in urine of patients that are BKVN positive per kidney allograft biopsy (BKVN), in urine collected in the first five days after transplant surgery (Early), urine collected from patients that are UTI negative per culture in the first month following transplantation (No UTI), samples collected before or after UTI (Pre-/Post-UTI) and samples collected at the time of UTI diagnosis (UTI). Two outliers in Pre-/Post-UTI and UTI groups correspond to the same patient, who suffered an acute rejection episode in the months prior. Low donor fractions in the Pre-/Post-UTI and UTI groups are likely due to increased immune cell presence in the urinary tract; patients with higher white blood cell counts have lower donor fractions (inset, red color indicates pyuria) **(b)** Absolute abundance of donor cfDNA in the urine of patients not diagnosed with infection in the first month post-transplant (red line is a LOESS filter smoothing curve, span = 1). Dotted lines connect samples from the same patient. **(c)** Genome coverage at the transcription start site (TSS), binned by gene expression level across all samples in the study. FPKM = Fragments per Kilobase of Transcript per Million mapped reads, an RNA-seq measure of gene expression. See supplemental table “Fig5A-B”.

Two studies recently demonstrated that the structure of chromatin in gene promoters is conserved within circulating cfDNA in plasma^22,21^. Ulz *et al*. employed whole-genome sequencing of plasma DNA to show that nucleosomal occupancy at transcription start sites results in different read depth coverage patterns for expressed and silent genes^22^. Here, we found that footprints of nucleosomes in gene promoters and transcriptional regulatory elements are conserved within urinary cfDNA (Fig. 5C, aggregation and normalization across all samples), and that the extent of nucleosomal protection is proportional to gene expression. Measurements of nucleosomal depletion can serve as a proxy for gene expression, and may be used to investigate host-pathogen interactions in the setting of UTI in more detail.

Mitochondrial DNA (mtDNA) in the urine was recently identified as a possible biomarker for hypertensive kidney damage^43^. Furthermore, recent data indicate a role for extracellular mitochondrial DNA as a powerful damage-associated molecular pattern (DAMP). Elevated levels of mtDNA in plasma have been reported in trauma, sepsis and cancer, and recent studies have identified mitochondrial DNA released into the circulation by necrotic cells^44^. For a small subset of patients diagnosed with BKVN (eight samples from seven subjects), we quantified donor and recipient specific mtDNA in urine, using an approach we have previously described^19^. We found that the graft is the predominant source of mitochondrial urinary cfDNA in seven of the eight samples (two-tailed Student t-test, *p* ≪ 10^−6^; see Methods). Molecular techniques to track DAMPs in urine released in the setting of kidney graft injury may provide a non-invasive window into the potential role of these molecules in the pathogenesis of immune-related complications.

## Discussion

We have presented a strategy to identify and assess infections of the urinary tract based on profiling of urinary cfDNA and ‘*omics* analysis principles. We show that different layers of clinical information are accessible from a single assay that are either inaccessible using current diagnostic protocols, or require parallel implementation of a multitude of different tests. In nearly all samples with clinically reported viral or bacterial infection of the urinary tract, cfDNA identified the suspected causative agent of infection. In addition, cfDNA sequencing revealed the frequent occurrence of cfDNA from bacteria that remain undetected in current clinical practice. In many samples, including those from patients regarded as clinically stable, we detected cfDNA from viruses that may be clinically relevant but not routinely assayed in the screening protocol at our institution. The assay we present therefore has the potential to become a valuable tool to monitor bacteriuria and viruria in transplant cohorts, and to ascertain their potential impact on allograft health.

Beyond measurement of the abundance of different components of the microbiome, urinary cfDNA provides a wealth of information about bacterial phenotypes. We show, for the first time, that analyses of the structure of microbial genomes from cfDNA allow estimation of bacterial population growth rates, thereby providing information about dynamics from a single snapshot. We compared the bacterial growth rates in samples with clinically-diagnosed UTI to those without diagnosed UTI and we observed higher growth rates for clinically-reported bacteria in patients diagnosed with UTI. We further show that metagenomic analysis of urinary cfDNA can be used to infer susceptibility to antimicrobials. We mined shotgun sequencing data for AR genes, and found a good agreement between the presence of AR genes and *in vitro* phenotypic antimicrobial susceptibility testing of bacterial isolates. cfDNA resistome profiling may have added potential over conventional antimicrobial resistance testing methods, as these methods typically use one or a few cultured colonies. cfDNA profiling can potentially capture AR gene fragments from the entire bacterial population which may be particularly important since cfDNA profiling revealed frequent putative co-infections within the UTI group.

Several new methodologies have been introduced in recent years to characterize the urinary microbiome and to diagnose urinary tract infection, including 16S ribosomal DNA sequencing^34,45,46^, and expanded culture techniques^8,47^. These approaches have challenged the clinical dogma that urine from healthy individuals is sterile^33^, and have revealed deficiencies in the culture protocols that are used in clinical practice today^9,34^. The cfDNA shotgun sequencing assay described here provides a versatile alternative that will be particularly useful for the monitoring of kidney transplant recipients, given the potential to enable viral and bacterial pathogen detection, antimicrobial resistance profiling, and graft injury monitoring from a single assay.

More than 15,000 patients receive lifesaving kidney transplants in the US each year^48^. Viral and bacterial infections of the urinary tract occur frequently in this patient group and often lead to serious complications, including graft loss and death. In the general population, UTI is one of the most frequent medical problems that patients present with in medical offices^49^. Shotgun sequencing of urinary cfDNA offers a comprehensive window into infections of the urinary tract and can be a valuable future diagnostic tool to monitor and diagnose bacterial and viral infections in kidney transplantation as well as in the general population. The assay we have presented is compatible with a short assay turnaround time (1-2 days), and will benefit from continued technical advances in DNA sequencing that will reduce cost and increase throughput in years to come.

## Methods

### Study cohort and sample collection

141 urine samples were collected from kidney transplant recipients who received care at New York Presbyterian Hospital – Weill Cornell Medical Center. We assayed urine samples from a total of 82 patients. We included 31 subjects who developed bacterial UTIs diagnosed within the first 12 months of transplantation and 14 subjects who never developed urinary tract infections within the first 3 months of transplantation. For the 31 subjects who developed UTIs, we assayed 43 urine samples corresponding to same day positive urine cultures (UTI Group); we assayed 15 urine samples from 15 subjects, collected at least 2 to 16 days (median 7 days) prior to development of the positive urine cultures (Pre-UTI Group), and we assayed 12 urine samples from 9 subjects, collected at least 3 to 26 days (median 9 days) after development of the positive urine cultures (Post-UTI Group) (7 of the 9 subjects were treated with antibiotics). We assayed a total of 29 samples collected within three months after transplantation from 14 subjects who never developed UTI in the first three months of transplantation. Ninety samples in the study had a corresponding same day urine culture with the associated urine specimens that were assayed. The study also included 25 samples from 23 subjects who had a corresponding positive diagnosis of BK virus nephropathy by needle biopsy of the kidney allograft (BKVN positive group) and 11 samples from 11 subjects who had a normal protocol biopsy and was negative for BK virus (BKVN negative group). Finally, the study additionally had 7 samples from 3 subjects who developed either clinically diagnosed rare viral infections including parvovirus or adenovirus. See also detailed clinical metadata in supplemental table “Clinical Data”.

### Urine collection and supernatant isolation

Most urine samples were collected via the conventional mid-stream void method (n=130). Samples obtained prior to post-transplant day 4 were collected via Foley catheter (n = 11). Approximately 50 mL of urine was centrifuged at 3,000 *g* for 30 minutes and the supernatant was stored at −80 °C in 1 or 4 ml aliquots. cfDNA was extracted from 1 mL (131 samples) or 4 mL (10 samples) of urine (Qiagen Circulating Nucleic Acid Kit, Qiagen, Valencia, CA).

### Analysis of discordance against bacterial culture

In a single sample, urinary cfDNA did not identify the uropathogen reported by conventional culture (*Raoultella ornithinolytica*). The patient had developed an *E. coli* UTI on post-operative day 6 and was treated initially with aztreonam but switched to cephalexin for a 14 day course. The subject subsequently developed a UTI that conventional bacterial culture revealed to be *R. ornithinolytica* on post-operative day 25. cfDNA analysis on the same day revealed a high abundance of *E. coli* UTI and no evidence of *R. ornithinolytica* infection. Given the discordant results, it is unclear if the second UTI is a recurrence as suggested by the cfDNA analysis or is a new infection as suggested by the urine culture data.

### Negative control

To control for environmental and sample-to-sample contamination, a known-template control sample (IDT-DNA synthetic oligo mix, lengths 25, 40, 55, 70 bp, 0.20 μM eluted in TE buffer) was included in every sample batch and sequenced to approximately 25% of the depth of the cfDNA extracts (~5 million fragments). The mean representation of each genus in the control was used to filter out genera in samples identified as possible contaminants. Possible sources of contamination in these experiments include: environmental contamination during sample collection in the clinic, nucleic acid contamination in reagents used for DNA isolation and library preparation, sample-to-sample contamination due to Illumina index switching^50^.

### Library preparation and next generation sequencing

Sequencing libraries were prepared using a single-stranded library preparation optimized for the analysis of ultrashort fragment DNA, described previously^19^. Libraries were characterized using the AATI fragment analyzer. Samples were pooled and sequenced on the Illumina NextSeq platform (paired-end, 2x75 bp). Approximately 50 million paired-end reads were generated per sample.

### Analysis – Composition of the urinary microbiome

Low quality bases and Illumina-specific sequences were trimmed (Trimmomatic-0.32^51^). Reads from short fragments were merged and a consensus sequence of the overlapping bases were determined using FLASH-1.2.7. Reads were aligned (Bowtie2, very sensitive mode^52^) against the human reference (UCSC hg19). Unaligned reads were extracted, and the non-redundant human genome coverage was calculated (SAMtools 0.1.19 rmdup^53^). To derive the urinary microbiome, reads were BLASTed (NCBI BLAST 2.2.28+) to a curated list of bacterial and viral reference genomes^54^. Short reads were assigned to specific taxa using a maximum likelihood algorithm that takes into account the ambiguity of read mapping^28,29,55^. The relative abundance of higher level taxa was determined on the basis of the genomic abundance at the strain or species level. For positive identification of viruses, we required at least 10 BLAST hits. In addition, due to the high load and genetic similarity of BK and JC polyomaviruses in many samples, we implemented a conservative filter for incompleteness and heterogeneity of genome coverage (GINI index less than 0.8 with at least 75% of the genome covered) for these two species only.

### Bacterial growth dynamics

Bacterial genome replication rates were determined using the methods described by Brown *et al.*^36^. Briefly, all bacterial strains within a sample were sorted and the GC-skew was used to identify the origin and terminus of replication (minimum and maximum GC-skew, respectively). Bacterial genomes were binned in 1 kbp tiles. The coverage was smoothed based on a running mean of 100 nearest neighboring tiles. The coverage in each tile was quantified and tiles were sorted by coverage. Linear regression was performed between the origin and terminus of replication after further removing the 5% least and most covered bins. The product of the slope of the regression line and the genome length was defined as the growth rate, a metric applied in previous analyses^36^. This analysis was applied for all bacterial strains with genome lengths greater than 0.5 Mbp, R^2^ linear regression correlation greater than 0.90, and GINI index coefficient less than 0.2, for which at least 2500 BLAST hits were detected in the sample.

### Nucleosome footprints in gene bodies

Paired-end reads were aligned using BWA-mem. The sequence read coverage in 2 kbp windows around the transcription start site was determined using the SAMtools depth function^22^.

### Proportion of donor-specific cfDNA in urine

The fraction of donor specific cfDNA in urine and plasma samples was estimated for sex-mismatched, donor-recipient pairs. The donor fraction was determined as follows:

Male donor, female recipient: *D* = 2*Y*/*A*,

Female donor, male recipient: *D* = 1-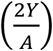,

where *Y* and *A* are the coverage of the mappability-adjusted Y and autosomal chromosomes, respectively. Sequence mappability was determined using HMMcopy^56^.

### Mitochondrial donor fraction

The proportion of donor specific mitochondrial DNA was quantified using methods previously described^19^. Briefly, a mitochondrial consensus sequence was determined for the donor and recipient from pre-transplant whole blood samples. Comparison of cfDNA sequence data to the mitochondrial consensus was used to quantify donor and recipient specific mitochondrial DNA. One sample was removed due to low depth of sequencing.

### Clinical antimicrobial resistance determination

Antimicrobial susceptibility testing was performed on 43 matched samples from patients with clinically diagnosed UTIs at New-York Presbyterian Hospital-Weill Cornell Medical Center. Antimicrobial susceptibility testing was organism-specific and included a combination of disk diffusion, gradient diffusion and microbroth dilution prepared and analyzed according to either the manufacturer’s or Clinical and Laboratory Standards Institute recommendations.

### Determining antimicrobial resistance gene presence

Nonhuman sequencing reads were aligned to a database of protein sequences of known antimicrobial resistance genes (CARD, 2,158 genes) using blastx (identity overlap 90%, culling limit 8 blastx hits). The hits with the highest identity and overlap length were selected for each read and compared to the antimicrobial resistance classes using the CARD ontology^38^.

### Statistical analysis

All statistical analyses were performed using R version 3.3.2. Unless otherwise noted, groups were compared using the nonparametric Mann-Whitney U test. Fourier analyses were performed using the spec.pgram function, part of the standard stats package, in R.

### Boxplots

Boxes in the boxplots indicate the 25^th^ and 75^th^ percentiles, the band in the box indicates the median, lower whiskers extend from the hinge to the smallest value at most 1.5 * IQR of the hinge, higher whiskers extend from the hinge to the highest value at most 1.5 * IQR of the hinge.

## Data availability

The sequencing data that support the findings of this study are made available in the database of Genotypes and Phenotypes (dbGaP).

## Acknowledgments

This work was supported by R21AI133331 (I.D.V. and J.R.L.), R21AI124237 (I.D.V.), DP2AI138242 (I.D.V.), K23AI124464 (J.R.L.), R37AI051652 (M.S.), and the Robert Noyce Foundation (I.D.V.). P.B. is supported by a NSF GRFP, DGE-1144153. We thank Erin Berthelsen for providing samples for assay development, and Catherine Snopkowski and Carol Li for help with sample handling and qPCR experiments.

## Materials & Correspondence

All requests should be submitted to I.D.V. and J.R.L.

## Author contributions

P.B., J.R.L, D.D., M.S. and I.D.V. contributed to the study design. P.B., M.H. and F.C. performed the experiments. P.B. J.R.L., D.D., L.F.W. and I.D.V. analyzed the data. P.B, D.D., L.F.W., J.R.L., and I.D.V. wrote the manuscript. All authors provided comments and edits.

## References

1. Foxman, B. Urinary tract infection syndromes. Occurrence, recurrence, bacteriology, risk factors, and disease burden. Infectious Disease Clinics of North America 28, 1–13 (2014).

2. Abbott, K. C. et al. Late urinary tract infection after renal transplantation in the United States. Am. J. Kidney Dis. 44, 353–362 (2004).

3. Ariza-Heredia, E. J. et al. Impact of urinary tract infection on allograft function after kidney transplantation. Clin. Transplant. 28, 683–690 (2014).

4. Chuang, P., Parikh, C. R. & Langone, A. Urinary tract infections after renal transplantation: a retrospective review at two US transplant centers. Clin. Transplant. 19, 230–235 (2005).

5. Hirsch, H. H. et al. Polyomavirus-associated nephropathy in renal transplantation: interdisciplinary analyses and recommendations. Transplantation 79, 1277–1286 (2005).

6. Dadhania, D. et al. Epidemiology of BK virus in renal allograft recipients: independent risk factors for BK virus replication. Transplantation 86, 521–528 (2008).

7. Schmiemann, G., Kniehl, E., Gebhardt, K., Matejczyk, M. M. & Hummers-Pradier, E. The Diagnosis of Urinary Tract Infection: A Systematic Review. Deutsches Ärzteblatt International 107, 361–367 (2010).

8. Price, T. K. et al. The Clinical Urine Culture: Enhanced Techniques Improve Detection of Clinically Relevant Microorganisms. J. Clin. Microbiol. 54, 1216–22 (2016).

9. Hilt, E. E. et al. Urine Is Not Sterile: Use of Enhanced Urine Culture Techniques To Detect Resident Bacterial Flora in the Adult Female Bladder. J. Clin. Microbiol. 52, 871–876 (2014).

10. Quake, S. Sizing up cell-free DNA. Clinical chemistry 58, 489–490 (2012).

11. Fan, H. C. et al. Non-invasive prenatal measurement of the fetal genome. Nature 487, 320–324 (2012).

12. Lo, Y. M. et al. Presence of fetal DNA in maternal plasma and serum. Lancet (London, England) 350, 485–487 (1997).

13. Tsui, N. B. Y. et al. High Resolution Size Analysis of Fetal DNA in the Urine of Pregnant Women by Paired-End Massively Parallel Sequencing. PLoS One 7, 1–7 (2012).

14. Fan, H. C., Blumenfeld, Y. J., Chitkara, U., Hudgins, L. & Quake, S. R. Noninvasive diagnosis of fetal aneuploidy by shotgun sequencing DNA from maternal blood. Proc. Natl. Acad. Sci. U. S. A. 105, 16266–16271 (2008).

15. De Vlaminck, I. et al. Circulating cell-free DNA enables noninvasive diagnosis of heart transplant rejection. Sci. Transl. Med. 6, 241ra77 (2014).

16. Bettegowda, C. et al. Detection of circulating tumor DNA in early- and late-stage human malignancies. Sci. Transl. Med. 6, 224ra24 (2014).

17. Meyer, M. et al. A High-Coverage Genome Sequence from an Archaic Denisovan Individual. Science 338, 222–226 (2012).

18. Gansauge, M.-T. & Meyer, M. Single-stranded DNA library preparation for the sequencing of ancient or damaged DNA. Nat. Protoc. 8, 737–48 (2013).

19. Burnham, P. et al. Single-stranded DNA library preparation uncovers the origin and diversity of ultrashort cell-free DNA in plasma. Sci. Rep. 6, 27859 (2016).

20. Korem, T. et al. Growth dynamics of gut microbiota in health and disease inferred from single metagenomic samples. Science 349, 1101–1106 (2015).

21. Snyder, M. W., Kircher, M., Hill, A. J., Daza, R. M. & Shendure, J. Cell-free DNA Comprises an In Vivo Nucleosome Footprint that Informs Its Tissues-Of-Origin. Cell 164, 57–68 (2016).

22. Ulz, P. et al. Inferring expressed genes by whole-genome sequencing of plasma DNA. Nat. Genet. 48, 1273–1278 (2016).

23. Botezatu, I. et al. Genetic analysis of DNA excreted in urine: a new approach for detecting specific genomic DNA sequences from cells dying in an organism. Clin. Chem. 46, 1078–1084 (2000).

24. Zhang, J. et al. Presence of donor-and recipient-derived DNA in cell-free urine samples of renal transplantation recipients: urinary DNA chimerism. Clin. Chem. 45, 1741–1746 (1999).

25. Su, Y.-H. et al. Human urine contains small, 150 to 250 nucleotide-sized, soluble DNA derived from the circulation and may be useful in the detection of colorectal cancer. J. Mol. Diagn. 6, 101–107 (2004).

26. Lo, Y. M. D. et al. Maternal Plasma DNA Sequencing Reveals the Genome-Wide Genetic and Mutational Profile of the Fetus. Sci. Transl. Med. 2, 61ra91–LP-61ra91 (2010).

27. Varshavsky, A. J., Bakayev, V. V, Chumackov, P. M. & Georgiev, G. P. Minichromosome of simian virus 40: presence of histone HI. Nucleic Acids Research 3, 2101–2113 (1976).

28. De Vlaminck, I. et al. Noninvasive monitoring of infection and rejection after lung transplantation. Proc. Natl. Acad. Sci. 112, 13336–13341 (2015).

29. Xia, L. C., Cram, J. A., Chen, T., Fuhrman, J. A. & Sun, F. Accurate genome relative abundance estimation based on shotgun metagenomic reads. PLoS One 6, (2011).

30. Ding, R. et al. CD103 mRNA levels in urinary cells predict acute rejection of renal allografts. Transplantation 75, 1307–1312 (2003).

31. Dadhania, D. et al. Validation of Noninvasive Diagnosis of BK Virus Nephropathy and Identification of Prognostic Biomarkers. Transplantation 90, 189–197 (2010).

32. Jm, R. Haemophilus influenzae pyelonephritis in adults. Arch. Intern. Med. 159, 316 (1999).

33. Wolfe, A. J. & Brubaker, L. ‘Sterile Urine’ and the Presence of Bacteria. Eur. Urol. 68, 173–174 (2015).

34. Wolfe, A. J. et al. Evidence of uncultivated bacteria in the adult female bladder. J. Clin. Microbiol. 50, 1376–1383 (2012).

35. Chaban, B. et al. Characterization of the vaginal microbiota of healthy Canadian women through the menstrual cycle. Microbiome 2, 23 (2014).

36. Brown, C. T., Olm, M. R., Thomas, B. C. & Banfield, J. F. Measurement of bacterial replication rates in microbial communities. Nat Biotech 34, 1256–1263 (2016).

37. Mollerup, S. et al. Propionibacterium acnes: Disease-Causing Agent or Common Contaminant? Detection in Diverse Patient Samples by Next-Generation Sequencing. J. Clin. Microbiol. 54, 980–987 (2016).

38. McArthur, A. G. et al. The Comprehensive Antibiotic Resistance Database. Antimicrobial Agents and Chemotherapy 57, 3348–3357 (2013).

39. Grskovic, M. et al. Validation of a Clinical-Grade Assay to Measure Donor-Derived Cell-Free DNA in Solid Organ Transplant Recipients. J. Mol. Diagn. 18, 890–902 (2016).

40. Lo, Y. D. et al. Presence of donor-specific DNA in plasma of kidney and liver-transplant recipients. Lancet 351, 1329–1330 (1998).

41. Smith, R. M. Urinary Infection in Children. N. Engl. J. Med. 205, 181–185 (1931).

42. Beck, J. et al. Digital droplet PCR for rapid quantification of donor DNA in the circulation of transplant recipients as a potential universal biomarker of graft injury. Clin. Chem. 59, 1732–1741 (2013).

43. Eirin, A. et al. Urinary Mitochondrial DNA Copy Number Identifies Chronic Renal Injury in Hypertensive Patients. Hypertens. (Dallas, Tex. 1979) 68, 401–410 (2016).

44. Lood, C. et al. Neutrophil extracellular traps enriched in oxidized mitochondrial DNA are interferogenic and contribute to lupus-like disease. Nat. Med. 22, 146–153 (2016).

45. Fouts, D. E. et al. Integrated next-generation sequencing of 16S rDNA and metaproteomics differentiate the healthy urine microbiome from asymptomatic bacteriuria in neuropathic bladder associated with spinal cord injury. J. Transl. Med. 10, 174 (2012).

46. Nelson, D. E. et al. Characteristic male urine microbiomes associate with asymptomatic sexually transmitted infection. PLoS One 5, e14116 (2010).

47. Thomas-White, K. J. et al. Incontinence medication response relates to the female urinary microbiota. Int. Urogynecol. J. 27, 723–733 (2016).

48. Hart, A. et al. OPTN/SRTR 2015 Annual Data Report: Kidney. Am. J. Transplant. 17, 21–116 (2017).

49. Foxman, B., Barlow, R., D’Arcy, H., Gillespie, B. & Sobel, J. D. Urinary Tract Infection: Self-Reported Incidence and Associated Costs. Ann. Epidemiol. 10, 509–515 (2000).

50. Sinha, R. et al. Index Switching Causes ‘Spreading-Of-Signal’ Among Multiplexed Samples In Illumina HiSeq 4000 DNA Sequencing. bioRxiv (2017).

51. Bolger, A. M., Lohse, M. & Usadel, B. Trimmomatic: A flexible trimmer for Illumina sequence data. Bioinformatics 30, 2114–2120 (2014).

52. Langmead, B. & Salzberg, S. L. Fast gapped-read alignment with Bowtie 2. Nat Methods 9, 357–359 (2012).

53. Li, H. et al. The Sequence Alignment/Map format and SAMtools. Bioinformatics 25, 2078–2079 (2009).

54. Kent, W. J. BLAT--the BLAST-like alignment tool. Genome Res. 12, 656–664 (2002).

55. De Vlaminck, I. et al. Temporal response of the human virome to immunosuppression and antiviral therapy. Cell 155, 1178–87 (2013).

56. Ha, G. et al. Integrative analysis of genome-wide loss of heterozygosity and monoallelic expression at nucleotide resolution reveals disrupted pathways in triple-negative breast cancer. Genome Res. 22, 1995–2007 (2012).

